# Species associations and management intensity modulate the habitat niche breadth of forest birds

**DOI:** 10.1101/2020.12.16.423074

**Authors:** Marco Basile, Thomas Asbeck, João M. Cordeiro Pereira, Grzegorz Mikusiński, Ilse Storch

## Abstract

Species associations can have profound effects on the realized habitat niche of species, indicating that habitat structure alone cannot fully explain observed abundances. To account for this aspect of community organization in niche modelling, we developed multi-species abundance models, incorporating the local effect of potentially associated species, alongside with environmental ones, targeting mainly forest management intensity. We coupled it with a landscape-scale analysis to further examine the role of management intensity in modifying the habitat niche in connection with the landscape context. Using empirical data from the Black Forest in southern Germany, we focused on the forest bird assemblage and in particular on the cavity nester and canopy forager guilds. We included in the analysis species that co-occur and for which evidences suggest association is likely. Our findings show that the local effect of species associations can moderate the effects of management intensity. We also found that species express a larger habitat niche breadth in intensively managed forests, depending on the landscape context. Species associations may facilitate the utilization of a broader range of environmental conditions under intensive forest management, which benefits some species over others. Such network of associations may be a relevant factor in the effectiveness of conservation-oriented forest management.

## 1. Introduction

In forest ecosystems, trees are ‘biotic modifiers’ [1] that may alter the biotic conditions and modulate the access to resources for other species. For instance, trees can alter the amount of light that reaches lower forest layers and profoundly impact the plant species composing the herb layer [2] or change micro-morphological and chemical soil properties and greatly affect soil organisms [3]. Similarly, the occurrence and abundance of forest taxa such as bats, birds or insects depends on tree structures such as rot holes and cavities providing resources to them [4–6]. The importance of biotic modifiers highlights the fact that assemblages are formed as a product of both environmental filtering (i.e. local environmental conditions acting as filters for local species sorting) and species interactions [7]. Environmental filtering depends on the main vegetation type found in an area, which can be subject to spatial and temporal variation [8]. In contrast, species interactions depend on intra- and inter-specific competition for resources [9], along with other ecological processes such as predation [10] or facilitation interactions [11].

Species interactions are often assumed based on patterns of species co-occurrences, in relation to a baseline occurrence rate dependent on the environmental conditions [12]. Despite criticism to this approach [13], the inclusion of an ‘interaction’ component can indeed improve the outcome of ecological niche modeling [14]. In this context, however, ‘interactions’ are reduced to co-occurrences, which may be non-random and point at least towards species association. The effect of interacting species can modify ecological niches within populations, highlighting inter-individual differences [15], and within meta-populations, e.g. following the introduction of allochthonous species [16].

The concomitance of biotic and abiotic factors and processes shapes the ecological niche of species, described as the Hutchinsonian niche [17,18]. This is further modulated by anthropogenic influences that often elicit different local responses within species’ distribution ranges, frequently leading to niche contraction and population decline when anthropogenic disturbance is dominant [19]. In the case of forests, management can be a fundamental determinant of the ‘habitat niche’ of forest species [20]. The habitat niche includes ‘the multidimensional space containing the compositional and structural features of a species’ environment that induce its occupancy’ [21], and can include also types of food, nest sites and microclimate but is mostly referred to by forest structure [22]. However, landscape characteristics can contribute to alter the realized habitat niche, despite the habitat conditions. For instance, the degree of forest fragmentation can affect the ability of species to spill over into suboptimal habitat types, a density-dependent process also influenced by regional abundances [21]. This often results in niche contractions [21], as the overall breadth of the realized habitat niches is related to the ability of species to cope with landscape context [23]. Generally, an interplay between abiotic conditions, habitat structure, landscape context, and species co-occurrences ultimately shapes the realized niches of species, with consequences on populations and communities of forest organisms.

In this study, we combined species local abundances and habitat niche modelling, using local and landscape features, to understand how forest management intensity modifies the realized habitat niche of co-occurring and potentially associated species. Specifically, we aimed at answering two questions: 1) how do local effects of associated species abundances modify their realized habitat niche; and 2) how does forest management intensity modify the habitat niche of species in different landscape contexts.

Many forest species are negatively impacted by forest management intensity [24–26], but can still persist in situations where interspecific associations allows them to exploit different environmental spaces [27–29]. We relied on multi-species modelling to infer about associations [12] and restricted our analysis to cavity-nesting and canopy-foraging bird species, which allowed us to assume direct interspecific interactions for nesting and foraging sites during the breeding season. In this way, we could account for possible associations among species (as for research question 1), and for those species directly relying on structures which are influenced by management (as for research question 2).

The cavity nester bird guild is composed by primary cavity nesters, i.e. those bird species that excavate their own holes in trees, and secondary cavity nesters, i.e. those that use holes generated by natural processes or excavated by primary cavity nesters [30]. The substrate where the cavity is located triggers competitive interactions among species (e.g. Balestrieri et al., 2018; Pasinelli, 2007). Experimental evidence indicates that the supply of holes can limit the number of cavity nesters [30,33]. Forest management for timber production is considered one of the main causes of shortages in cavity supply [34], usually reflected by a decline of the cavity nester guild within the bird assemblage [35]. The main reason is that such management leads to the shortage of trees or snags with characteristics enabling formation of natural cavities or allowing their excavation by primary cavity nesters [36]. This is in contrast to primeval forests, where the abundance of cavities is not a limiting factor for cavity nesters and woodpeckers are not the key cavity providers [37]. Hence, in managed forests species-specific responses might show a high degree of variation depending on the relative abundance of large, suitable trees and snags [38,39], relative abundance of woodpeckers [40], forest management type [41], and tree species composition [42]. Moreover, the predation risk [43,44] and the presence of invasive species [45] are additional factors shaping the cavity nester guild.

The canopy forager guild includes those species that feed on substrates in the tree canopy [46]. In this case, competition sparks from the optimal foraging substrate (e.g. twigs vs. needles in conifer canopy) and the different efficiency of each species’ foraging technique [47,48]. In Europe, it comprises mainly foliage gleaners and seed eaters, and it is also influenced by forest management [49–51].

We modelled the habitat niches of co-occurring and potentially associated birds of the cavity nester and canopy forager guilds accounting for the habitat structure, the management intensity, the availability of tree-related microhabitats (TreMs) [52], and the landscape composition.

## 2. Methods

### 2.1 Study area

The study was performed in 127 one-hectare plots located in the forest landscape of the Black Forest, southwestern Germany, (Latitude: 47.6°- 48.3° N, Longitude: 7.7°-8.6° E; WGS 84). Plots were selected in the framework of the ConFoBi Research Training Group ([53]; confobi.uni-freiburg.de). Selection focused on stands with age over 60 years managed with continuous-cover forestry and located at least 1 km from each other. The plots ranged in terms of elevation between 500 and 1400 m a.s.l., and represented a typical temperate mixed mountain forest, dominated by Norway spruce (*Picea abies*), European beech (*Fagus sylvatica*) and silver fir (*Abies alba*).

### 2.2 Environmental predictors

The environmental descriptors included in the analysis were collected at the plot scale (1 ha), containing forest structure data, and at the landscape scale (up to 5 km²), including landscape configuration metrics and forest cover.

#### 2.2.1 Forest variables

The forest structure of the plots was described in terms of tree basal area, share of conifer trees, richness and abundance of TreMs [54] and an index of forest management intensity based [55], tree species composition and deadwood volume. An inventory used to describe forest structure comprised species identity and diameter at breast height (DBH) of all living trees (with DBH > 7 cm), from which basal area and the share of conifer trees were derived. In addition, the DBH of all snags (with DBH > 7 cm and height > 1.3 m) on the plots was measured and lying deadwood data was collected using the line intersect method [56]. The abundance and richness of TreMs was retrieved from previous research in the same plots [54]. TreMs are considered to be “a distinct, well delineated structure occurring on living or standing dead trees, that constitutes a particular and essential substrate or life site for species or species communities during at least a part of their life cycle to develop, feed on, using as shelter or to breed” [52] and have shown correlations to the richness and abundance of forest-dwelling vertebrates and (saproxylic) insects [4,57,58]. The forest management intensity index (ForMI) measured three different management aspects [55]: a) the proportion of harvested tree volume compared to the maximum volume; b) the proportion of species not belonging to the natural tree species composition; and c) the ratio of artificial (showing signs of cutting) vs. natural deadwood. The index spans values 0-3, where 0 would indicate a forest not managed for timber production and 3 an intensively managed production forest. In addition, the average altitude of each plot was provided from a digital terrain model with spatial resolution of 0.5 m (State Office for Geoinformation and Land Development Baden-Württemberg, Germany).

#### 2.2.2 Landscape variables

The landscape-scale predictors included the forest cover-based metrics describing the fragmentation of the forest surrounding the plots. Forest cover was derived from the land cover map provided by the State Office for Geoinformation and Land Development of Baden-Württemberg, Germany (Geobasdata ©, www.lgl-bw.de, ref no: 2851.9-1/19). Forest cover was assessed in the neighboring five km^2^ circular area, separately for conifer and total forest. The landscape metrics were computed on the same land cover map, using only areas classified as forest for the computation (i.e. binary map), and employed the software FRAGSTATS [59]. We considered six metrics commonly employed to describe fragmentation and patchiness of the landscape: the aggregation index, the contiguity area index, the core area index, the edge density, the Euclidean nearest neighbor and the landscape shape index. These metrics were selected because either evidence or experts suggest they have an effect on the numerical response of birds [60].

### 2.3 Bird sampling

Birds were sampled using standardized point counts with limited distance of 50 m, repeated up to three times/year during the period March-June in years 2017-2019 (i.e. encompassing most of the breeding season), starting half an hour after sunrise with the latest sample collected at 12:00 CET. Each survey lasted 20 minutes, during which birds were recorded repeatedly for every 5 minutes, in order to reach a reasonable sample coverage [61,62].

Each bird species was classified as cavity nester, insectivorous canopy forager, or both according to standard references [30,63,64]. Species recorded less than 30 times in the entire study were excluded from the analysis. To estimate the effect of the co-occurrence and abundance of a given species on a potentially associated one, we established linkages between species, i.e. assumed that the abundance of a species has an effect on the abundance of another one. To inform the direction of the linkage, i.e. whether species A has an effect on species B or the opposite, we relied on the existing literature (see appendix S1).

### 2.4 Abundance estimates

Bird abundance was estimated using community N-mixture models [65,66]. Such models allow for estimation of abundance of the species belonging to an animal assemblage as a function of environmental predictors, taking into account the detectability error by employing repeated count data. Our models incorporated the density-dependent effect of co-occurring and potentially associated species (appendix S2). Species were paired according to their relationship (as described in appendix S1), and assumptions about the effect were not made. That is, after establishing from the literature that the species A has an effect on species B, we did not assume this effect to be positive or negative, but only density-dependent. We restricted our analysis by focusing only on the cavity nester and canopy forager species found in our study area, which potentially compete over resources. Hence, if a relationship was present, the abundance of species A was considered a covariate of the response variable, i.e. the abundance of the species B, similarly to other research on species co-occurrences [67]. Species abundance was modelled as a Poisson process, while the detectability was modelled as a binomial distribution, dependent on the abundance process and moderated by the date and time of the surveys to model individual heterogeneity in detectability. The forest variables included in the species abundance model had a variance inflation factor ≤ 3, indicating that multicollinearity was not an issue. All predictors were scaled prior to analysis. The full model was built in JAGS programming language and fitted by applying Bayesian inference. We used uninformative priors and ran three MCMC chains of 400 000 iterations, discarding the first 10 000 and thinning by 90. We considered reliable model parameter estimates those drawn from a posterior distribution where the proportion with the same sign as the mean was *f* ≥ 0.9. If this was not the case, the model parameter was discarded from the analysis and the model ran again. We considered that chains reached convergence when the Gelman-Rubin statistic (r-hat) was ≤ 1.1 for all parameters [68]. All analyses were conducted in R environment. The community model was built with the package ‘jagsUI’ [69].

### 2.5 Habitat niche characterization

The forest structure of each plot was characterized by performing hierarchical clustering of the forest variables, aimed at grouping them in two clusters of high management intensity (HMI) and low management intensity (LMI) plots. Each plot was assigned to a category of management intensity using the K-means clustering method on the forest variables. At the landscape scale, instead, we characterized the landscape structure by building two gradients using the first two components of a Principal Component Analysis (PCA) on the correlation matrix of landscape variables. Considering that species perceive the landscape according to their life histories, we performed PCAs on the guilds and on each species. Since more or less fragmented landscape configurations can result in very different bird abundances [70,71], we considered optimal landscapes as those where the estimated abundance at plot level was higher than the mean abundance observed across the study area (0, when scaled). In this way, we characterized both the landscape with high (HA) and low (HA) abundances, as proxies for optimal and suboptimal-to-unsuitable landscapes, and each included both HMI and LMI plots. We performed a permutational Multivariate ANalysis Of VAriance (perMANOVA), to test whether each landscape variable derived in the neighborhood of plots differed among management intensity classes to further confirm that landscape structure was independent from forest structure. Once the forest and landscape structure were characterized, we compared the habitat niche breadth of the guilds and each species both when accounting for species associations, and by projecting it only as a function of forest variables. Secondly, we compared the habitat niche breadth for HMI and LMI plots along the landscape gradients in LA and HA landscapes. We calculated the abundance of each guild by summing up the respective species’ estimated abundances and scaling them. The habitat niche breadth was visualized by plotting the 95% confidence ellipses of abundance estimates and visually comparing the respective niche position and width. The R package ‘vegan’ was employed for the analysis [72].

## 3. Results

### 3.1 Guilds abundance and associations

Over three years, we recorded 8812 individuals belonging to 16 different species that were detected at least 30 times, of which were 11 cavity nesters, 10 canopy foragers, with 5 belonging to both guilds (table 1). The most common species counted at plots was the coal tit (*Periparus ater*), with an average number of individuals per plot and visit of 0.93 (± 0.84 SD), while the short-toed treecreeper (*Certhia brachydactyla*) was the least common (0.02 ± 0.17 SD). Multi-species model included 77 associations between the 16 species (figure S1). Species deemed not to be influenced by other species included woodpeckers, large-sized species (e.g. stock dove *Columba oenas*), species that can substantially rely on other resources (e.g. blackcap *Sylvia atricapilla* forage also in shrubs), or species that can escape competition by various means (e.g. European nuthatch *Sitta europaea* can substantially modify the cavity opening). The mean probability of detection across all species was 0.18, ranging from 0.03 (± 0.69 SD) of the blackcap to 0.51 (± 0.55 SD) of the great tit (*Parus major*). All 16 species responded in abundance to at least one forest variable, reporting a credible estimate (*f* < 0.9), while for ten species the effect of the forest management intensity index (ForMI) scored a credible and negative estimate (table 1). Co-occurring species scored credible association estimates with up to four other species, totaling between the two guilds 22 credible associations, out of the 77 hypothesized (figure 1). Statistical associations indicated thirteen negative and nine positive effects between two species. Three species, the great tit, the marsh tit (*Poecile palustris*), and the short-toed treecreeper, returned only one positive association, while four associations were found for the blue tit (*Cyanistes caeruleus)*. More associations were identified among canopy foragers than among cavity nesters.

**Table 1:**
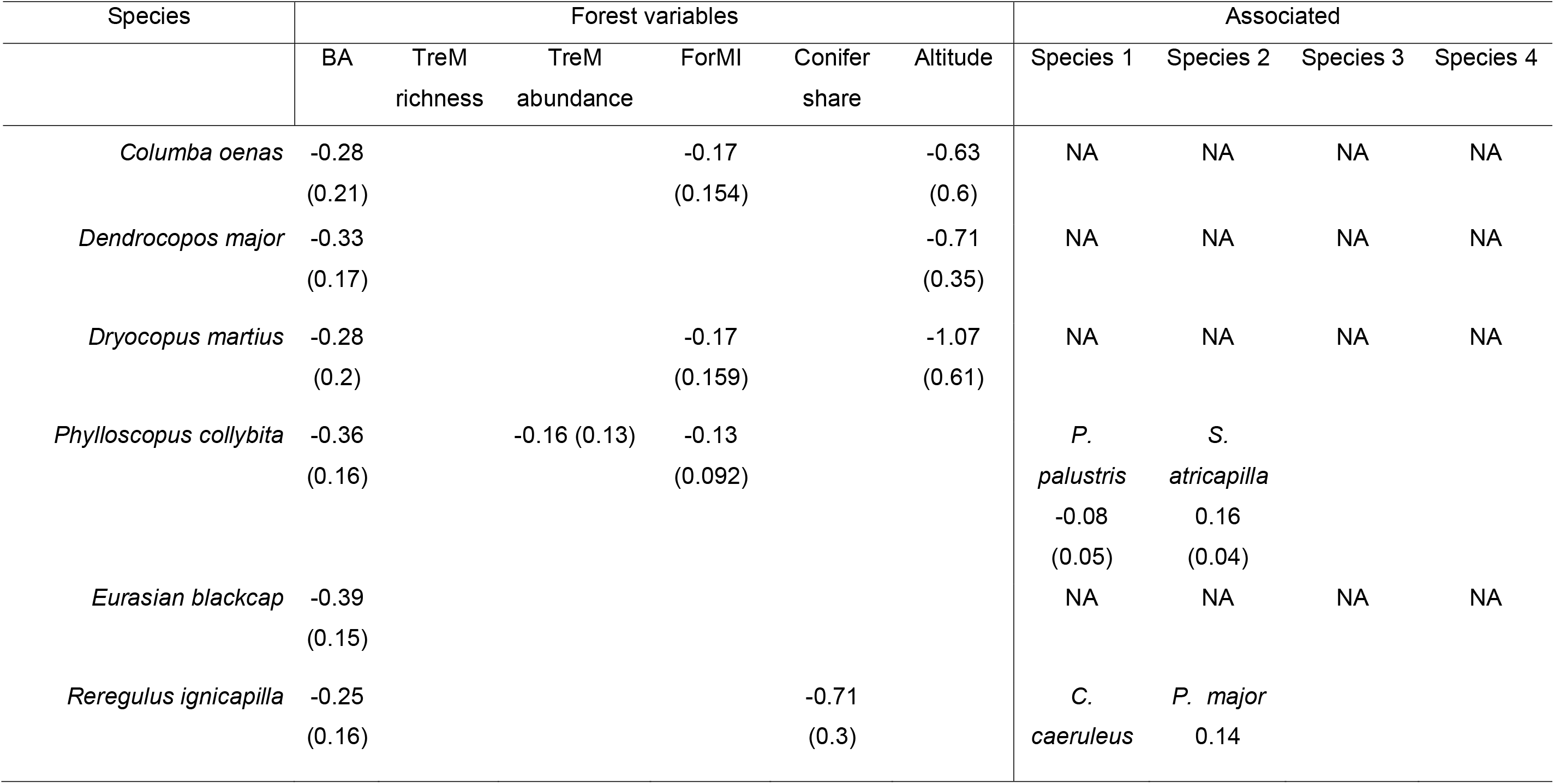
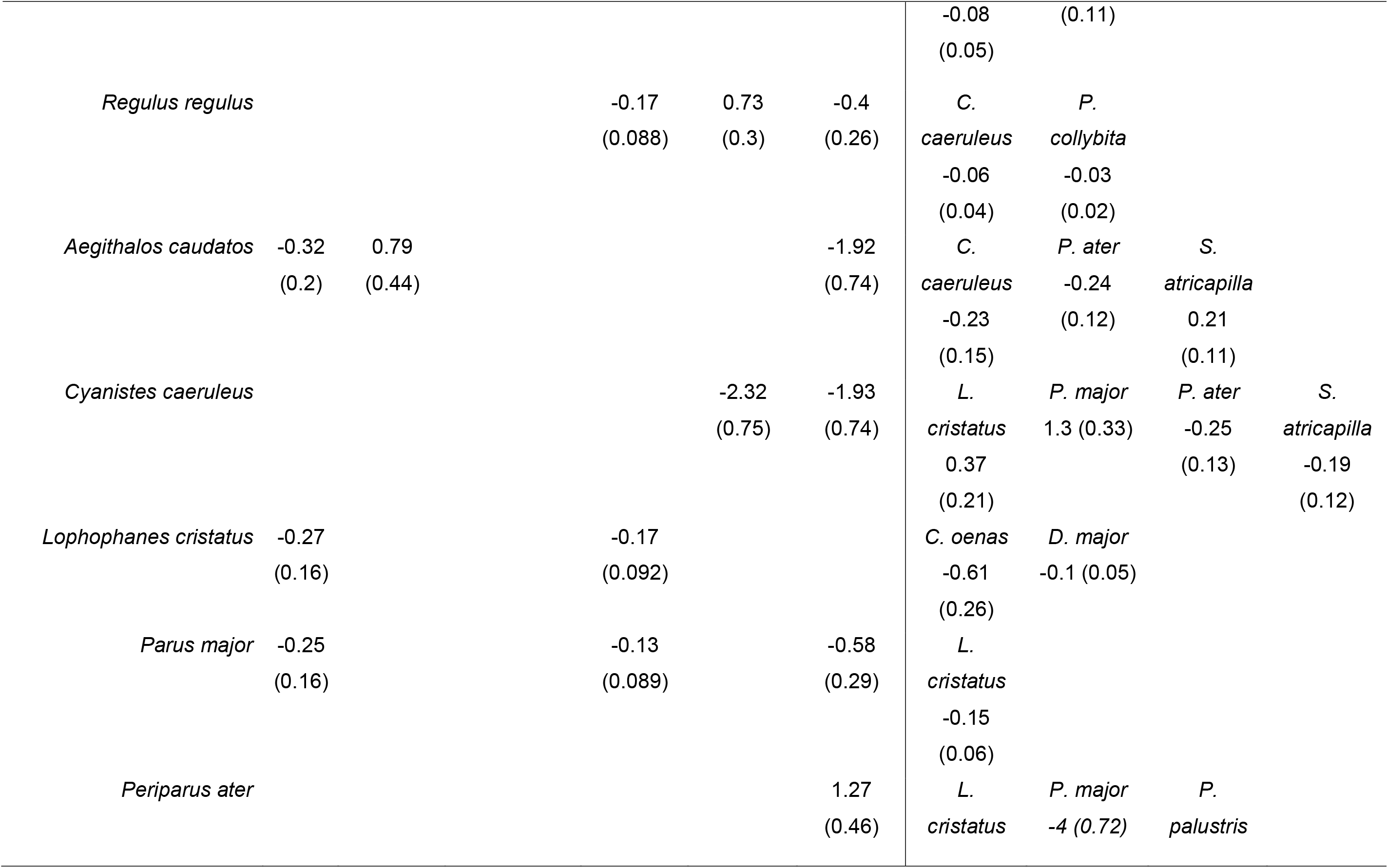
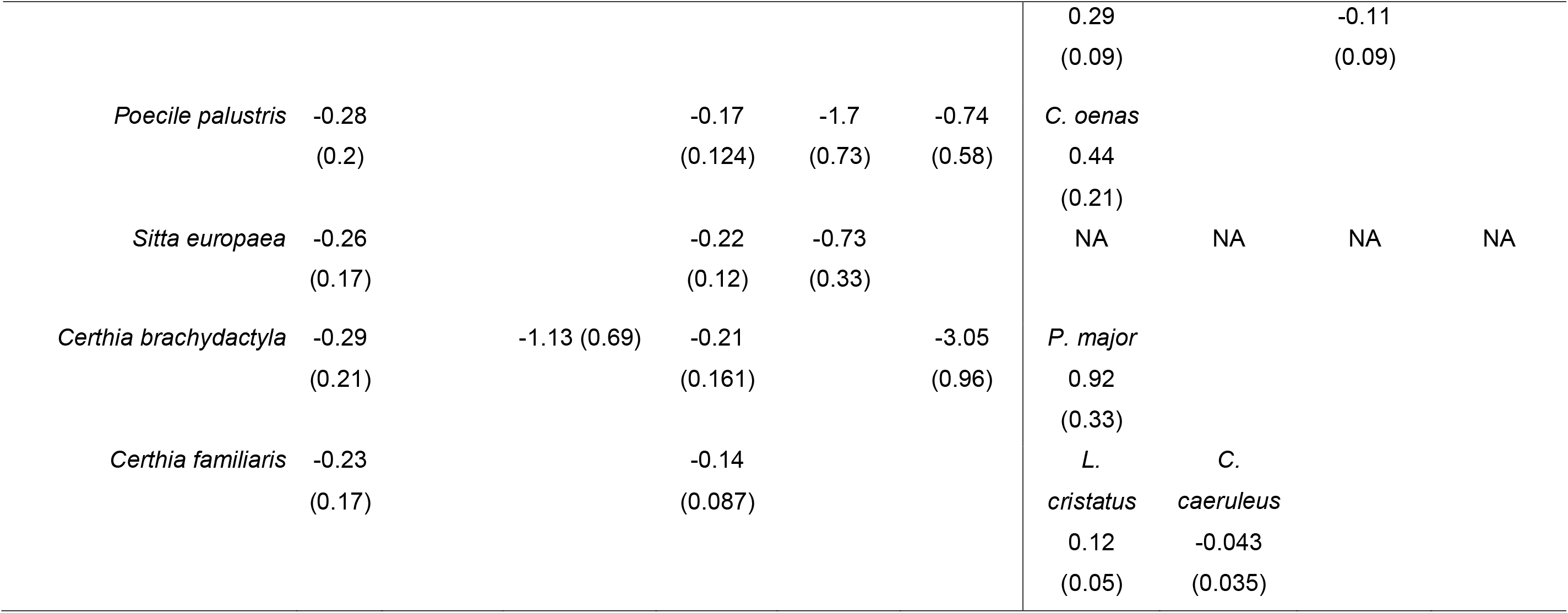
Effect estimates (with standard deviations) of environmental predictors and associated species on species abundances considered in the study belonging to either the cavity nester or canopy forager guild. BA = basal area; ForMI = forest management intensity index; NA = not applicable to species considered not to be influenced by other species due to life history traits (see methods and Appendix S1).

**Figure 1:**
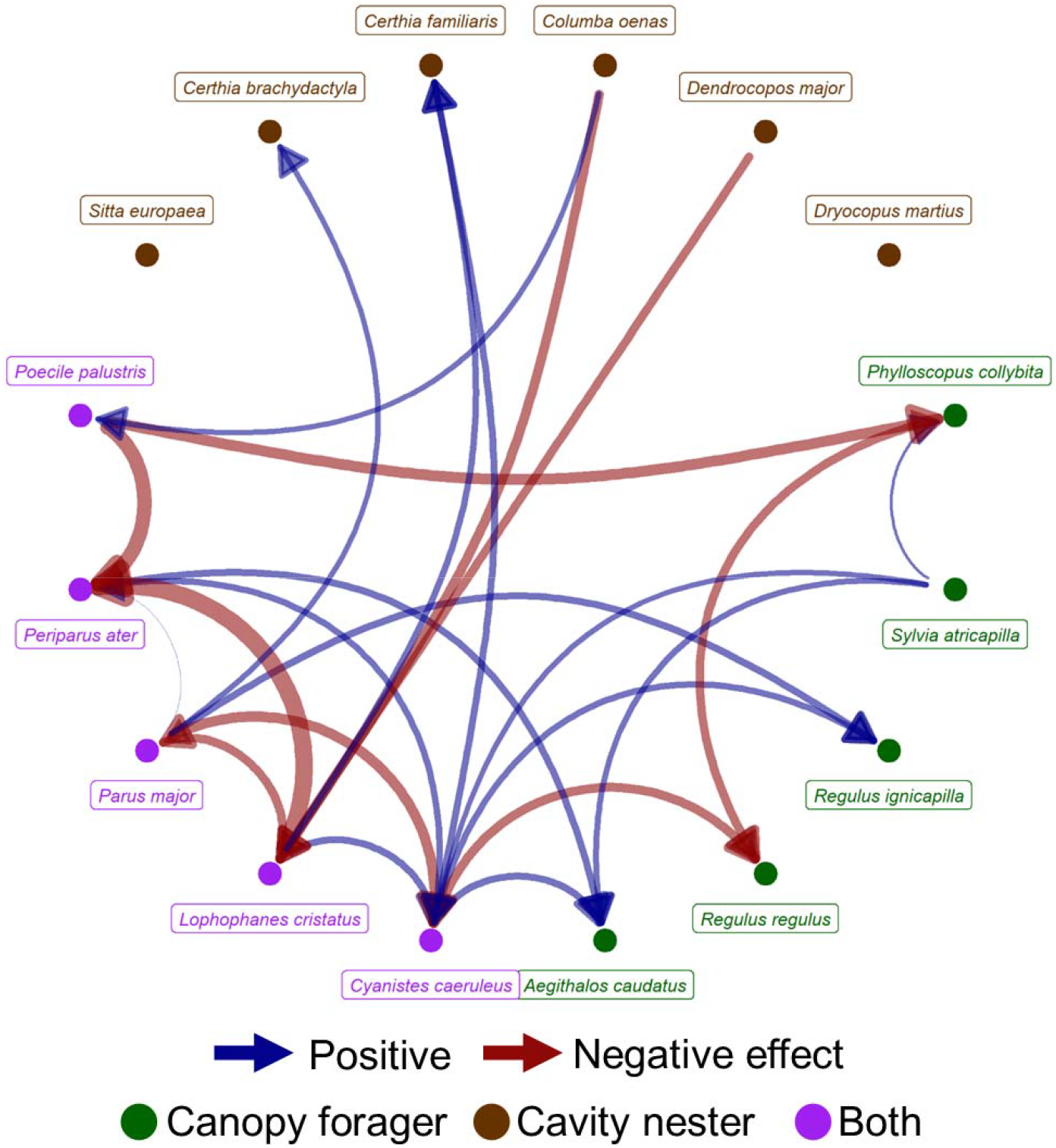
Statistical associations among species found by modelling species abundances as a function of co-occurring species and environmental predictors. Twenty-two positive and negative associations were found. Associations are considered as density-dependent effects on the abundance of the influenced species, possibly due to interspecific competition over resources, in this case, nesting and feeding resources. Arrow width indicates the effect size.

### 3.2 Habitat niche characterization

Based on the forest variables identified as abundance predictors, the hierarchical clustering identified 79 plots as belonging to high management intensity calss (HMI) plots, with means of basal area = 35.66 (± 7.87 SD), conifer share = 0.76 (± 0.20 SD), ForMI = 1.47 (± 0.36 SD), TreM richness = 8.04 (± 2.72 SD), and TreM abundance = 30.38 (± 14.39 SD). Low management intensity (LMI) plots were 48 with means of basal area = 37.88 (± 12.64 SD), conifer share = 0.51 (± 0.25 SD), ForMI = 0.78 (± 0.44 SD), TreM richness = 14.5 (± 4.49 SD), and TreM abundance = 67.4 (± 48.41 SD). Altitude was similar between management intensity classes. Landscape variables were not correlated with management intensity classes, with all perMANOVAs scoring p > 0.05.

For the entire studied assemblage, we did not find differences in the breadth of the habitat niche along the gradients of forest variables, when including species associations in the habitat niche model (figure 2). Along the altitudinal gradient the niche breadth was also similar in both models, although the model without associations presented a rather positive response, due to the high abundances estimated for the coal tit. However, absolute abundances per species and plot were mostly larger in the model accounting for associations: higher abundances are spread along the ForMI gradient (figure 3). This was particularly evident for the crested tit (*Lophophanes cristatus*) and the Eurasian treecreeper (*Certhia familiaris*), despite both having negative associations with other species.

**Figure 2:**
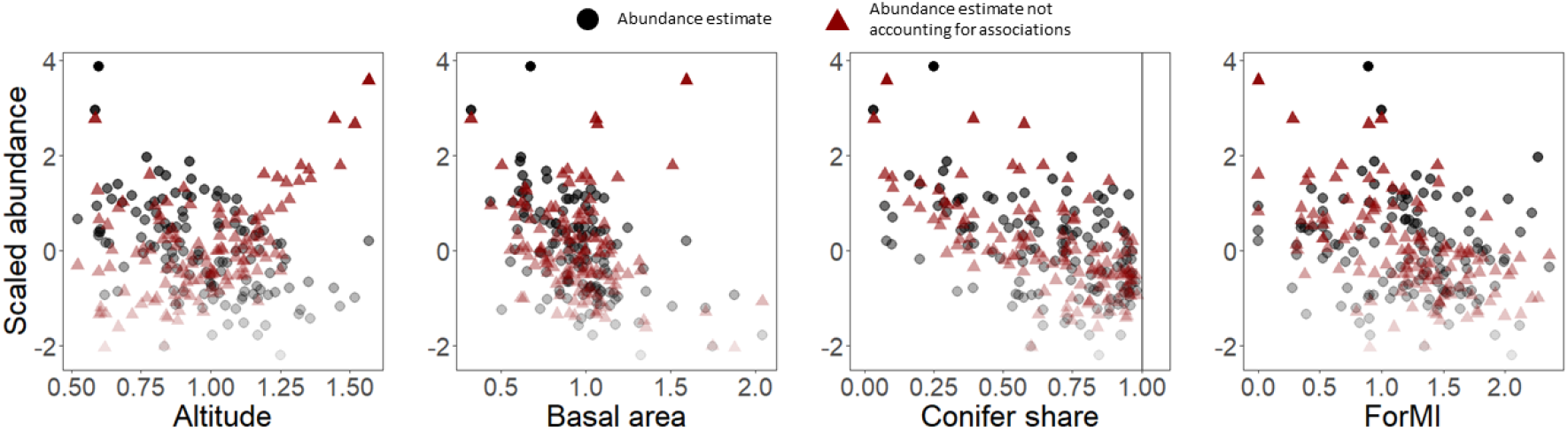
Scaled abundance of the bird assemblage including cavity nester and canopy forager guilds estimated from 127 forest plots in the Black Forest, Germany. Red triangles show estimates as a function of environmental predictors, while black circles show estimates incorporating also the statistical associations among species. Transparency denotes abundance values.

**Figure 3:**
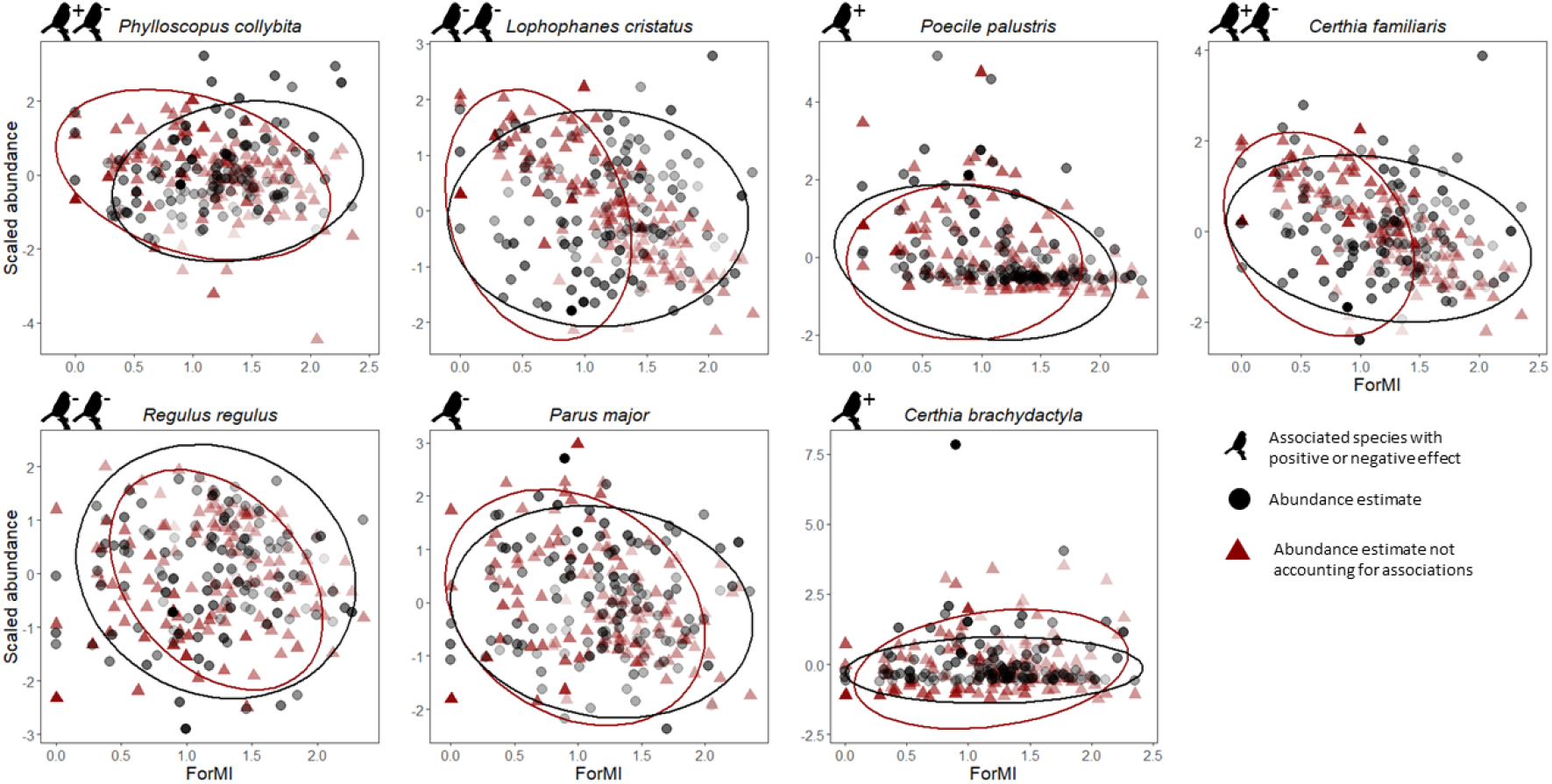
Scaled abundance of the species influenced by forest management intensity index (ForMI). Red triangles show estimates as a function of environmental predictors, while black circles show estimates incorporating also the statistical associations among species. 95% ellipses coded with same colors encircle the section along the environmental gradient where the upper quartile of the abundances occur. Bird icons indicate the number of associated species with positive (+) and negative (-) effects.

The PCA of the landscape variables for high abundance (HA) landscapes of the entire bird assemblage could explain ~ 63 % of the variation observed. The first component scored positive loadings for aggregation index, forest and conifer forest cover, and negative loadings for edge density and landscape shape index. This suggests that high abundances are observed in landscapes with large, contiguous forest patches (table 2). Low abundance (LA) landscapes differed in the loadings of contiguity index, core area, forest and conifer forest cover. This indicates low abundances located in landscapes with less forest cover and smaller forest patches. In this case the variance explained was ~ 61 % (table 2). The bird assemblage did not differ in habitat niche breadth between HMI and LMI plots, in HA landscapes. In contrast, in LA landscapes the habitat niche was smaller in LMI plots than in HMI plots. This indicates that the assemblage widened the habitat niche in more fragmented landscapes under high management intensity (figure 4). A similar pattern was observed for the single species within the cavity nester and canopy forager guilds, especially among the associated ones (figures 5–7). Specifically, Eurasian treecreeper, chiffchaff (*Phylloscopus collybita*), European blackcap, firecrest (*Regulus ignicapilla*), goldcrest (*Regulus regulus*), blue tit, coal tit exhibited a larger niche breadth in HMI plots located in LA landscapes. Among the other species, stock dove and marsh tit showed an opposite pattern, with larger niche breath in HMI plots from HA landscapes. The woodpeckers, short-toed treecreeper, great tit, and crested tit had niche breadths larger in HMI plots for both landscape categories. Finally, the niche breadth of European nuthatch and long-tailed tit (*Aegithalos caudatus*) were similar in all landscapes.

**Table 2:** Factor loadings of the first two components (PC) of the principal component analysis on the landscape variables for high abundance (HA) and low abundance (LA) landscapes. Variance explained by each component is in brackets

| Landscape variable | HA landscape |  | LA landscape |  |
| --- | --- | --- | --- | --- |
|  | PC1 | PC2 | PC1 | PC2 |
|  | (44.43 %) | (18.20 %) | (46.29 %) | (14.93 %) |
| Aggregation index | 0.49 | 0.06 | 0.46 | 0.30 |
| Contiguity index | 0.01 | -0.68 | 0.12 | -0.41 |
| Core area | 0.26 | -0.53 | 0.31 | 0.08 |
| Edge density | -0.41 | 0.09 | -0.43 | -0.25 |
| Euclidean nearest neighbour | -0.17 | 0.41 | -0.16 | 0.43 |
| Landscape shape index | -0.50 | -0.07 | -0.47 | -0.27 |
| Forest cover | 0.32 | 0.23 | 0.29 | -0.56 |
| Conifer forest cover | 0.37 | 0.11 | 0.41 | -0.30 |

**Figure 4:**
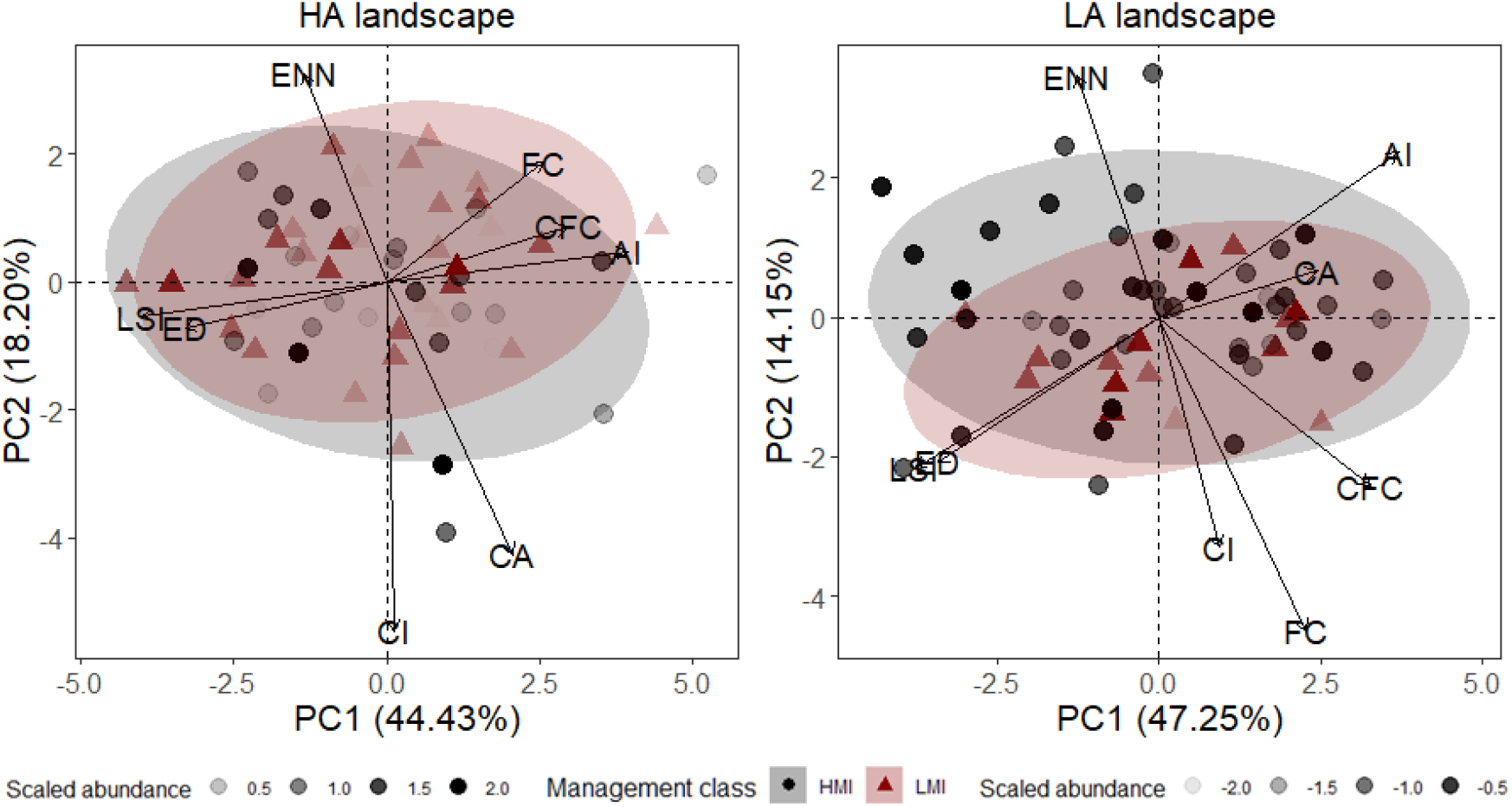
Habitat niche breadth of the bird assemblage of the Black Forest, Germany, including cavity nesters and canopy foragers, visualized as 95% confidence ellipses from PCA of landscape variables, for high (HA) and low (LA) abundance landscapes, respectively. High management intensity (HMI) and low management intensity (LMI) plots are determined by a hierarchical clustering on forest variables. Numbers on axis represent the variance explained by the principal component. AI = aggregation index; CA = core area; CI = contiguity index; CFC = conifer forest cover; ED = edge density; ENN = Euclidean nearest neighbour; FC = forest cover; LSI = landscape shape index.

**Figure 5:**
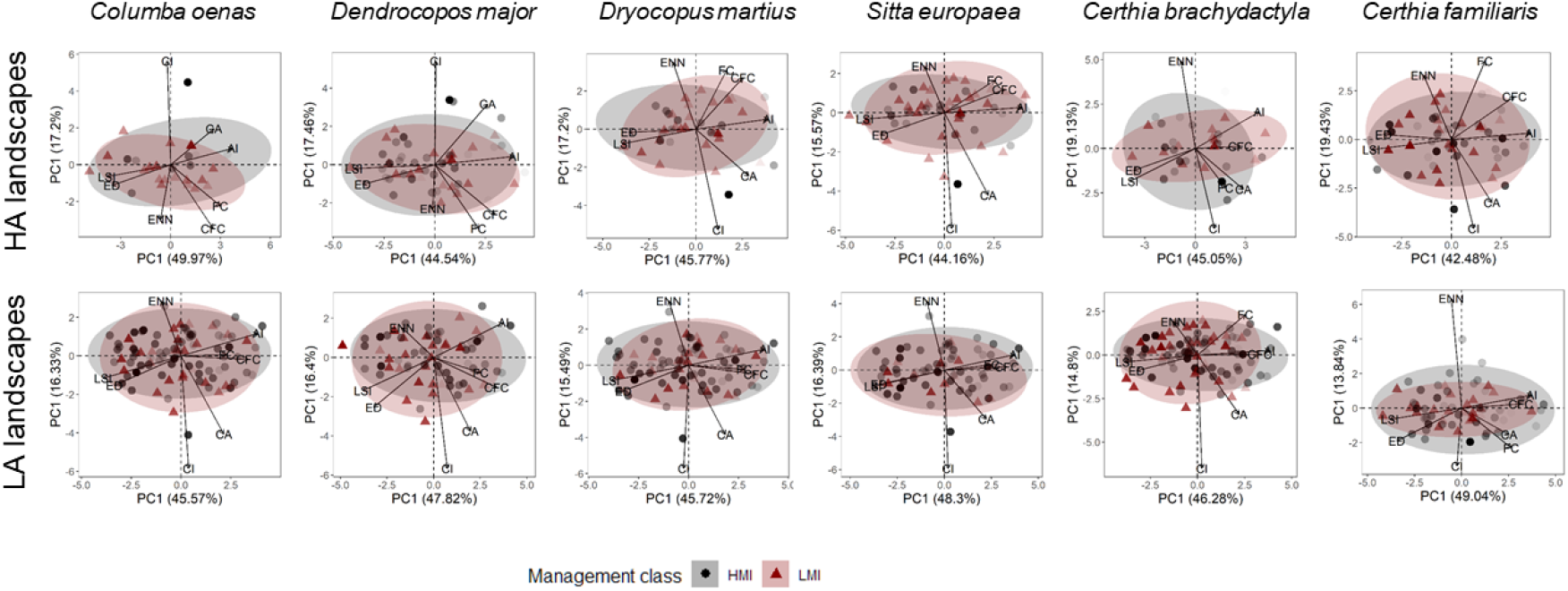
Habitat niche breadth of cavity nester species (excluding canopy foragers), visualized as 95% confidence ellipses from PCA of landscape variables, for high (HA) and low (LA) abundance landscapes, respectively. High management intensity (HMI) and low management intensity (LMI) plots are determined by a hierarchical clustering on forest variables. Numbers on axis represent the variance explained by the principal component. AI = aggregation index; CA = core area; CI = contiguity index; CFC = conifer forest cover; ED = edge density; ENN = Euclidean nearest neighbour; FC = forest cover; LSI = landscape shape index. From left to right, species are stock dove, great spotted woodpecker, black woodpecker, European nuthatch, short-toed treecreeper, Eurasian treecreeper.

**Figure 6:**
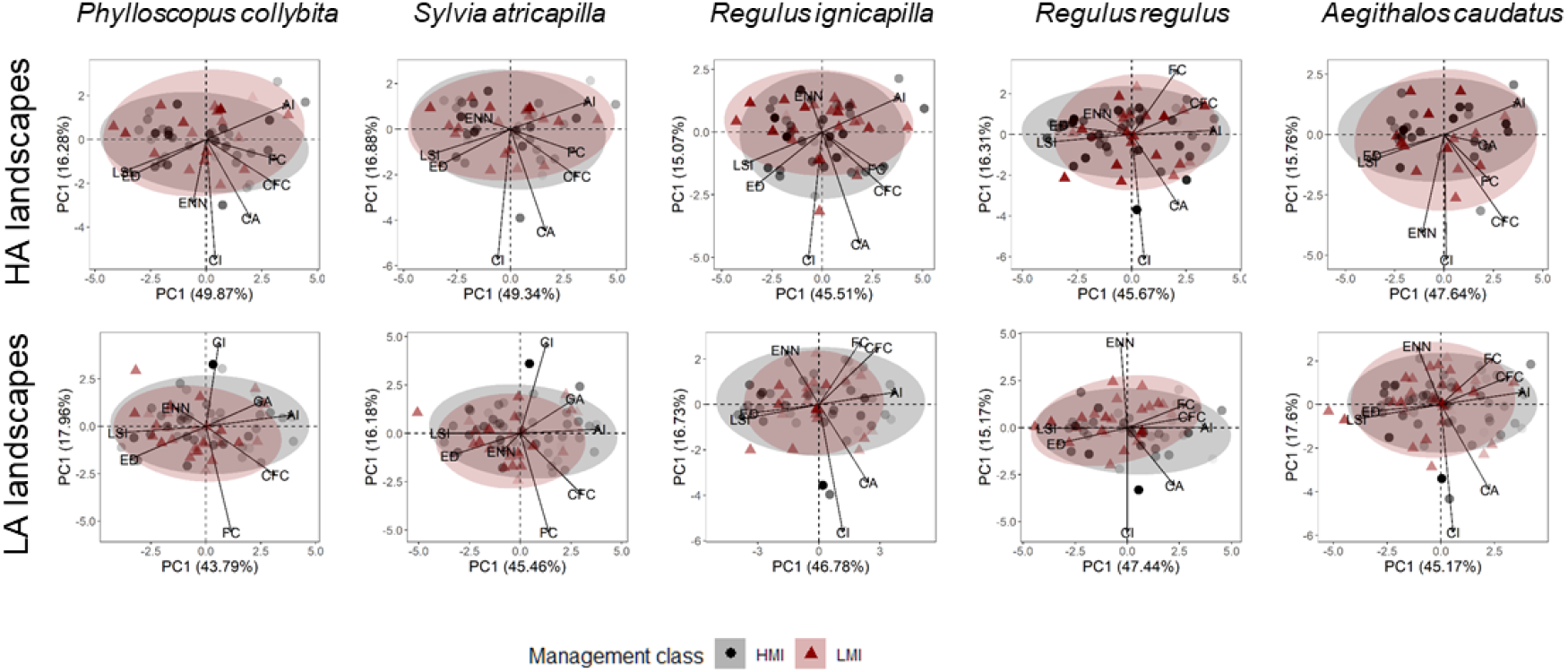
Habitat niche breadth of canopy forager species (excluding cavity nesters), visualized as 95% confidence ellipses from PCA of landscape variables, for high (HA) and low (LA) abundance landscapes, respectively. High management intensity (HMI) and low management intensity (LMI) plots are determined by a hierarchical clustering on forest variables. Numbers on axis represent the variance explained by the principal component. AI = aggregation index; CA = core area; CI = contiguity index; CFC = conifer forest cover; ED = edge density; ENN = Euclidean nearest neighbour; FC = forest cover; LSI = landscape shape index. From left to right, species are chiffchaff, European blackcap, firecrest, goldcrest, long-tailed tit.

**Figure 7:**
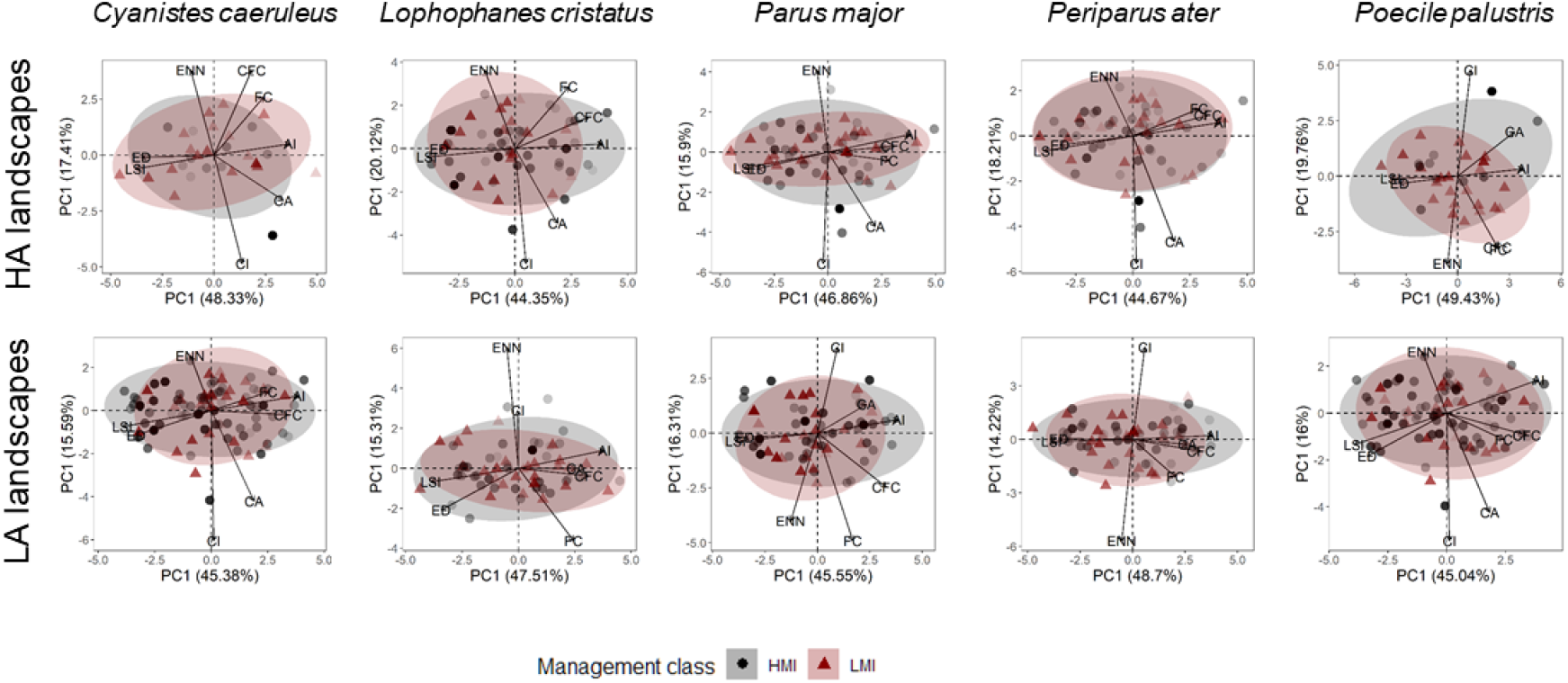
Habitat niche breadth of species belonging to both cavity nester and canopy forager guilds, visualized as 95% confidence ellipses from PCA of landscape variables, for high (HA) and low (LA) abundance landscapes, respectively. High management intensity (HMI) and low management intensity (LMI) plots are determined by a hierarchical clustering on forest variables. Numbers on axis represent the variance explained by the principal component. AI = aggregation index; CA = core area; CI = contiguity index; CFC = conifer forest cover; ED = edge density; ENN = Euclidean nearest neighbour; FC = forest cover; LSI = landscape shape index. From left to right, species are blue tit, crested tit, great tit, coal tit, marsh tit.

## 4. Discussion

### 4.1 Species associations and the realized habitat niche of co-occurring species

The role of species associations in modulating the habitat niche depends on the context in which they occur (i.e. the abiotic, spatial, temporal and community context), which may alter their magnitude and direction [73]. Large-scale environmental drivers have previously been linked with alterations in species association and co-occurrence patterns [74]. In our study, using forest birds, we demonstrated that the occurrence and abundance of species at a small (plot-level) spatial scale, are linked to local environmental drivers and, simultaneously, to the associated species. Hence, we observed a context-dependent change in the realized habitat niche of species. Considering the Eurasian treecreeper as an example, based on our results, we would predict lower abundances at sites with high management intensity. However, the positive association between the crested tit and the Eurasian treecreeper predicts the latter to still persist in higher abundances at intensively managed sites. The mechanism behind the observed process that increasing numbers of crested tits is associated with an increasing abundance of the Eurasian treecreeper roots likely in the network of relationships within the entire bird assemblage. However, we did not directly observe that, but rather interpreted statistical associations as direct or indirect influence of a species on another one, causing the abundance of the species being influenced to change according to density-dependent processes, e.g. competition over resources [75].

Most of the species association effects were less important than forest variable effects, indicating that the primary determinant of species abundance remains the habitat structure. However, habitat structure cannot fully explain the processes of habitat selection and community assembly, without considering it alongside with other processes, such as competition or predation [7,76]. Compelling evidence shows that it is not possible to separate the effect of environmental filtering from that of biotic interactions in traditional study settings [8]. Indeed, we show that the abundances of associated species (a proxy for biotic interactions) are related to each other and modulate the response of species to environmental gradients (environmental filtering). This raises the question whether associations play a similarly important role within forest bird guilds other than those investigated whenever the set of resources they use is limited by forest management.

The effect of species associations sheds new light on the effect of forest variables on the targeted bird species. Conifer share at the plot scale influenced abundances of species mainly in a negative way, agreeing with other studies from the same [77] and other regions [78]. The effects of basal area and ForMI were mainly negative among species, despite different degrees of forest specialization among species. Indeed, species responded negatively to management intensity, which is common among cavity nesters [50,79]. Instead, forest generalist species, such as the great tit, are often associated with increasing management intensity [80]. Nonetheless, we observed a negative response to management intensity for this species. Our model also showed that this negative response is softened by species associations, observing high abundances in high management intensity plots. The great tit is often found in coexistence with other members of the family Paridae, which may result in mutualistic relationships [81,82]. This coexistence is based on different types of resource partitioning leading to unique combinations of niche characteristics [47,75]. Hence, we stress that, regarding cavity nesters, positive effects on abundance linked with high management intensity may be an artifact of complex patterns of co-occurrence and association among species rather than a direct consequence of the habitat structure.

### 4.2 Management intensity modifies the habitat niche according to the landscape context

Birds can exhibit larger territory areas, lower occupancy rates, and lower abundances in suboptimal habitats [70,83–85]. In landscapes characterized by low abundances of forest birds (LA landscapes in our study), species may then be forced to exploit a broader range of environmental conditions. Our analysis showed that birds occurred in high management intensity (HMI) plots across a wider set of environmental conditions (at the landscape scale), especially in suboptimal landscapes (i.e. with lower abundances). In contrast, we found that when abundances are higher across the entire landscape (regardless of local conditions), the habitat niches of a given species from differently managed plots tend to overlap. This is in agreement with a previous studies relating higher occurrence rates with more generalist habits in birds [23,86]. Another study focusing on boreal forest birds provided findings similar to ours, indicating that local habitat structure is a more important driver of habitat niche breadth rather than landscape structure [20]. The major determinant of habitat niche breadth in our study was management intensity. Species with and without associations showed similar patterns across all landscapes, and the responses among species were consistent with each other. Therefore, we can provide support to the statement that management intensity can modify the habitat niche of forest birds, with the remark that this process is more evident in suboptimal landscapes. Low abundances may be a result of fragmentation, sub-optimal habitat quality, disturbance, or other factors [87]. In this study, low abundances are negatively associated with forest patch size and forest cover. In fact, the landscape context may act as precondition for a species to utilize different environments [88,89]. Moreover, studies focusing only on foraging guilds and including old-growth forests (barely represented in our study) associated wider niche breadths with more pristine forests [90,91]; this is partly in accordance with our results in HA landscapes and LMI plots for Eurasian treecreeper, chiffchaff and long-tailed tit. This showcases a great difficulty in optimizing conservation interventions for the widest possible array of species in forest management strategies targeting the conservation of forest bird species, and especially cavity nesters, the guild dominated in Europe by resident species with many of them being in jeopardy in managed forests [92–94]. It has been demonstrated that forest management may, through reducing niche diversity, largely affects the ability of closely-related species and competing species to co-exist [76].

### 5 Conclusions

Interspecific associations can result in striking differences in habitat selection for bird species with almost completely overlapping habitat niches [95,96]. This highlights the plasticity of birds in their habitat use, and in their ability to exploit different habitat conditions. We showed that within the forest bird assemblage, species can show different habitat niches according to the presence and abundance of co-occurring and associated species. Our approach could evaluate only unidirectional effects between species pairs, hence we could not test all possible association effects, which may be bidirectional in some instances [43]. This study suggests that interspecific associations can result in higher bird abundances in more intensively managed forests, which may be a consequence of factors beyond the local habitat structure. Hence, to design management actions based exclusively on species-habitat relationships may not deliver the best possible results, and managers should be aware of potential effects of interspecific associations. In addition, we did not include potential predation effects affecting the targeted species, an important density-dependent process that can potentially deliver different results [10,76]. However, many species within the guilds are similar in size, life history, and have mutualistic anti-predatory relationships [97]. At the same time, in suboptimal landscapes species are forced to exploit a broader range of environmental conditions in intensively managed forests, which may hamper the effectiveness of conservation-oriented management wherever landscape-scale factors are not taken into account. Forest managers need to develop flexible strategies that incorporate appropriate measures at the local scale, e.g. the retention of high-quality habitat trees offering suitable resources for cavity nesting birds [98], reducing the management intensity in this way.

## 6. Acknowledgements

Funding for this research was provided by the German Research Foundation (DFG), under the ConFoBi Research Training Group “Conservation of Forest Biodiversity in Multiple-Use Landscapes of Central Europe” (number GRK 2123). We are very thankful to J. Wildraut, C. Pacioni and M. von Vequel-Westernach for the support provided in the field collection of bird data, C. Pacioni for the help in managing the data, to F. Hauck and J. Grossmann for the support provided in the TreM and ForMI data collection, to J. Dilger, D. Saladin and J. Memmert for conducting the forest inventory.

