## Appendix S1 for "Species associations and management intensity modulate the habitat niche breadth of forest birds"

**Appendix S1.** Species associations and management intensity modulate the habitat niche breadth of forest birds.

Species classification in the cavity nester and canopy forager guilds. Based on the reference, we established 77 associations.

Cavity nester

*Primary cavity nester*

The primary cavity nesters include species that excavate their own holes in trees [1], namely the bird family Picidae (Vigors, 1825). The substrate where the cavity is formed triggers competitive interactions among species [2,3]. It is not rare that the same individuals use the same hole in multiple years, or that the same individuals excavate more than one hole per year, triggering competition among cavity nesters (including non-bird species) for the occupation of a given hole. The choice of the nesting hole is primarily driven by body size, given that some species are simply too large to fit in small cavities, while the quality of the nesting hole, such as the exposure to predation, triggers competition [4]. Woodpecker holes can last up to 10 years (median value) in coniferous forests, especially those excavated by large woodpecker species [5], while non-excavated holes have a shorter lifespan (median = 4.4 years) in coniferous forests [6]. The role of cavities in forests has been described using network theory, given the potentially large number of species connected through this key structure [7].

*Secondary cavity nester*

The secondary cavity nesters include species that use holes generated by natural processes or excavated by primary cavity nesters [1]. The availability of cavities in primeval forests is not a limiting factor for secondary cavity nesters [8]. Instead, in managed forests species-specific responses might show a high degree of variation depending on the predation risk [4,9], the relative abundance of large, suitable trees and snags [10,11], the relative abundance of woodpeckers [12], the forest management type [13], the tree species composition [14], and the presence of invasive species [15]. An abundant family of this guild is Paridae (Vigors, 1825), which includes small songbirds that also mainly forage in the canopy. The breeding ecology of this family, and especially the great and the blue tit (*Parus major* and *Cyanistes caeruleus*), have been the focus of vast literature. The larger species are considered dominant in the competition for the nesting hole, and, even though the smaller species can escape competition by relying on smaller holes, they can still suffer competition when cavities are limited [4,16,17]. The only secondary cavity nester excluded from the competition dynamic is the Eurasian nuthatch (*Sitta europaea*), given its ability to modify the entrance of a cavity [4].

Canopy forager

The bird canopy forager guild include those species which feeding substrate is found in the tree canopy [18]. In this case, competition sparks from the optimal foraging substrate-end (e.g. twigs vs. needles in conifer canopy) and the different efficiency of each species to forage [19,20]. In Europe, it comprises mainly foliage gleaners and seed eaters, and it is also influenced by forest management [21–23]. Other species included in this guild are large-sized species, such as the Eurasian jay (*Garrulus glandarius*). From the composition of this guild, we concluded that only foliage gleaners could have interaction with the cavity nesters over the nesting site or the feeding ground, hence we limited our analysis only to the insectivorous canopy forager guild. In this case, competition is driven by the ability of each species to cope with the local condition and the presence of other species, rather than by a limiting resource. Different part of the canopy (e.g. branches, twigs, or needles) or different forest layers offer multiple alternatives for feeding, with species having partly overlapping niches [24–27]. However, some species, such as the coal tit (*Periparus ater*) are particularly efficient at food exploitation, given some environmental condition, that can escape competition from larger species by food depletion [17,19,28]. Other species can also escape competition by feeding on the ground [29].

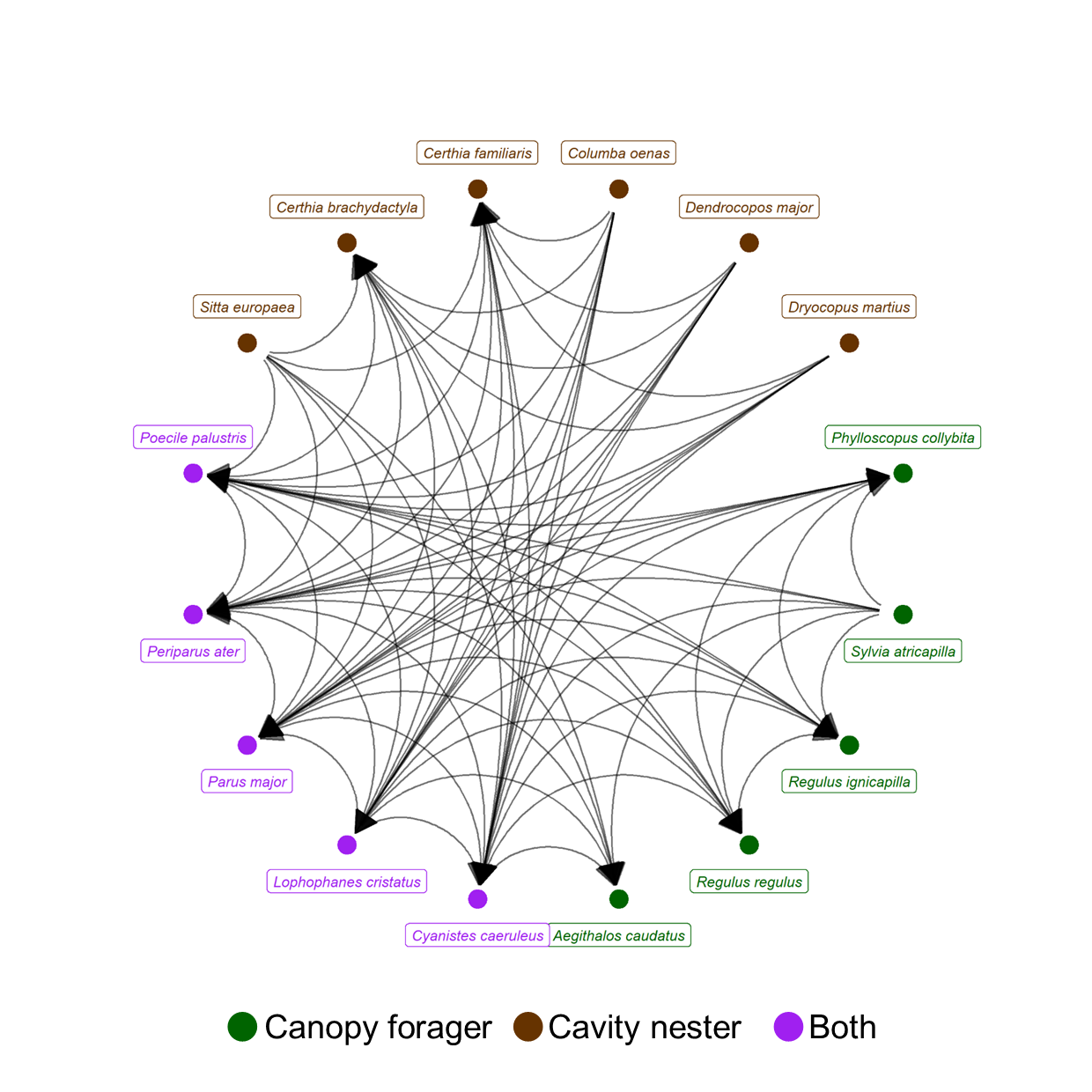

Figure S1: Relationship between species identified or suggested in the literature. The arrow points at the species that is influenced. The relationships have been included in the model described in S2. We interpret the relationships as influence on the abundance, due to competition over resources.
