## Appendix S2 for "Species associations and management intensity modulate the habitat niche breadth of forest birds"

**Appendix S2.** Species associations and management intensity modulate the habitat niche breadth of forest birds.

Multi-species abundance model incorporating species competition processes, based on a modified version of the community N-mixture models (Yamaura et al. 2012; Kéry & Royle 2016).

model {

\# Bird assemblage priors for species-specific parameters

for(k in 1:nspec){

alpha0[k] ~ dnorm(mu.alpha0, tau.alpha0)	# Detection intercepts

beta0[k] ~ dnorm(mu.beta0, tau.beta0)	# Abundance intercepts

for(v in 1:2){

alpha[k, v] ~ dnorm(mu.alpha[v], tau.alpha[v]) # Slopes detection

}

for(w in 1:6){

beta[k, w] ~ dnorm(mu.beta[w], tau.beta[w])	# Slopes abundance

}

}

\# abundance model

mu.beta0 ~ dunif(-2,2)

tau.beta0 <- pow(sd.beta0, -2)

sd.beta0 ~ dunif(0, 3)

for(w in 1:6){

mu.beta[w] ~ dunif(-2, 2)

tau.beta[w] <- pow(sd.beta[w], -2)

sd.beta[w] ~ dunif(0, 2)

}

\# Priors for interspecific association effects

CC_AC.phi ~ dnorm(0, 0.01) # blue tit vs long-tailed tit

CC_PC.phi ~ dnorm(0, 0.01) # blue tit vs chiffchaff

CC_RI.phi ~ dnorm(0, 0.01) # blue tit vs firecrest

CC_RR.phi ~ dnorm(0, 0.01) # blue tit vs goldcrest

CC_CB.phi ~ dnorm(0, 0.01) # blue tit vs short-toed treecreeper

CC_CF.phi ~ dnorm(0, 0.01) # blue tit vs treecreeper

DM_CC.phi ~ dnorm(0, 0.01) # great spotted woodpecker vs blue tit

DM_LC.phi ~ dnorm(0, 0.01) # great spotted woodpecker vs crested tit

DM_PM.phi ~ dnorm(0, 0.01) # great spotted woodpecker vs great tit

DM_PA.phi ~ dnorm(0, 0.01) # great spotted woodpecker vs coal tit

DM_CB.phi ~ dnorm(0, 0.01) # great spotted woodpecker vs short-toed treecreeper

DM_CF.phi ~ dnorm(0, 0.01) # great spotted woodpecker vs treecreeper

DM_PP.phi ~ dnorm(0, 0.01) # great spotted woodpecker vs marsh tit

LC_AC.phi ~ dnorm(0, 0.01) # crested tit vs long-tailed tit

LC_PC.phi ~ dnorm(0, 0.01) # crested tit vs chiffchaff

LC_RI.phi ~ dnorm(0, 0.01) # crested tit vs firecrest

LC_RR.phi ~ dnorm(0, 0.01) # crested tit vs goldcrest

LC_CB.phi ~ dnorm(0, 0.01) # crested tit vs short-toed treecreeper

LC_CF.phi ~ dnorm(0, 0.01) # crested tit vs treecreeper

LC_PP.phi ~ dnorm(0, 0.01) # crested tit vs marsh tit

LC_PA.phi ~ dnorm(0, 0.01) # crested tit vs coal tit

LC_CC.phi ~ dnorm(0, 0.01) # crested tit vs blue tit

LC_PM.phi ~ dnorm(0, 0.01) # crested tit vs great tit

PM_AC.phi ~ dnorm(0, 0.01) # great tit vs long-tailed tit

PM_PC.phi ~ dnorm(0, 0.01) # great tit vs chiffchaff

PM_RI.phi ~ dnorm(0, 0.01) # great tit vs firecrest

PM_RR.phi ~ dnorm(0, 0.01) # great tit vs goldcrest

PM_CC.phi ~ dnorm(0, 0.01) # great tit vs blue tit

PM_CB.phi ~ dnorm(0, 0.01) # great tit vs short-toed treecreeper

PM_CF.phi ~ dnorm(0, 0.01) # great tit vs treecreeper

PM_PP.phi ~ dnorm(0, 0.01) # great tit vs marsh tit

PA_AC.phi ~ dnorm(0, 0.01) # coal tit vs long-tailed tit

PA_PC.phi ~ dnorm(0, 0.01) # coal tit vs chiffchaff

PA_RI.phi ~ dnorm(0, 0.01) # coal tit vs firecrest

PA_RR.phi ~ dnorm(0, 0.01) # coal tit vs goldcrest

PA_CC.phi ~ dnorm(0, 0.01) # coal tit vs blue tit

PA_CB.phi ~ dnorm(0, 0.01) # coal tit vs short-toed treecreeper

PA_CF.phi ~ dnorm(0, 0.01) # coal tit vs treecreeper

PC_RI.phi ~ dnorm(0, 0.01) # chiffchaff vs firecrest

PC_RR.phi ~ dnorm(0, 0.01) # chiffchaff vs goldcrest

SA_AC.phi ~ dnorm(0, 0.01) # blackcap vs long-tailed tit

SA_CC.phi ~ dnorm(0, 0.01) # blackcap vs blue tit

SA_PM.phi ~ dnorm(0, 0.01) # blackcap vs great tit

SA_PC.phi ~ dnorm(0, 0.01) # blackcap vs chiffchaff

SA_RI.phi ~ dnorm(0, 0.01) # blackcap vs firecrest

SA_RR.phi ~ dnorm(0, 0.01) # blackcap vs goldcrest

SA_PP.phi ~ dnorm(0, 0.01) # blackcap vs marsh tit

SE_CC.phi ~ dnorm(0, 0.01) # nuthatch vs blue tit

SE_LC.phi ~ dnorm(0, 0.01) # nuthatch vs crested tit

SE_PM.phi ~ dnorm(0, 0.01) # nuthatch vs great tit

SE_PA.phi ~ dnorm(0, 0.01) # nuthatch vs coal tit

SE_CB.phi ~ dnorm(0, 0.01) # nuthatch vs short-toed treecreeper

SE_CF.phi ~ dnorm(0, 0.01) # nuthatch vs treecreeper

SE_PP.phi ~ dnorm(0, 0.01) # nuthatch vs marsh tit

CO_CC.phi ~ dnorm(0, 0.01) # stock dove vs blue tit

CO_LC.phi ~ dnorm(0, 0.01) # stock dove vs crested tit

CO_PM.phi ~ dnorm(0, 0.01) # stock dove vs great tit

CO_PA.phi ~ dnorm(0, 0.01) # stock dove vs coal tit

CO_CB.phi ~ dnorm(0, 0.01) # stock dove vs short-toed treecreeper

CO_CF.phi ~ dnorm(0, 0.01) # stock dove vs treecreeper

CO_PP.phi ~ dnorm(0, 0.01) # stock dove vs marsh tit

BW_CC.phi ~ dnorm(0, 0.01) # black woodpecker vs blue tit

BW_LC.phi ~ dnorm(0, 0.01) # black woodpecker vs crested tit

BW_PM.phi ~ dnorm(0, 0.01) # black woodpecker vs great tit

BW_PA.phi ~ dnorm(0, 0.01) # black woodpecker vs coal tit

BW_CB.phi ~ dnorm(0, 0.01) # black woodpecker vs short-toed treecreeper

BW_CF.phi ~ dnorm(0, 0.01) # black woodpecker vs treecreeper

BW_PP.phi ~ dnorm(0, 0.01) # black woodpecker vs marsh tit

PP_AC.phi ~ dnorm(0, 0.01) # marsh tit vs long-tailed tit

PP_PC.phi ~ dnorm(0, 0.01) # marsh tit vs chiffchaff

PP_RI.phi ~ dnorm(0, 0.01) # marsh tit vs firecrest

PP_RR.phi ~ dnorm(0, 0.01) # marsh tit vs goldcrest

PP_CB.phi ~ dnorm(0, 0.01) # marsh tit vs short-toed treecreeper

PP_CF.phi ~ dnorm(0, 0.01) # marsh tit vs treecreeper

PP_PA.phi ~ dnorm(0, 0.01) # marsh tit vs coal tit

PP_PM.phi ~ dnorm(0, 0.01) # marsh tit vs great tit

PP_CC.phi ~ dnorm(0, 0.01) # marsh tit vs blue tit

\# detection model

mu.alpha0 ~ dunif(-2, 2)

tau.alpha0 <- pow(sd.alpha0, -2)

sd.alpha0 ~ dunif(0, 2)

for(v in 1:2){

mu.alpha[v] ~ dunif(-2, 2)

tau.alpha[v] <- pow(sd.alpha[v], -2)

}

sd.alpha[1] ~ dunif(0, 2)

sd.alpha[2] ~ dunif(0, 2)

\# Ecological model for true abundance (process model)

for (i in 1:nsite){

N[i,1] ~ dpois(lambda[i,1]) # long-tailed tit

N[i,2] ~ dpois(lambda[i,2]) # blue tit

N[i,3] ~ dpois(lambda[i,3]) # great spotted woodpecker

N[i,4] ~ dpois(lambda[i,4]) # crested tit

N[i,5] ~ dpois(lambda[i,5]) # great tit

N[i,6] ~ dpois(lambda[i,6]) # coal tit

N[i,7] ~ dpois(lambda[i,7]) # chiffchaff

N[i,8] ~ dpois(lambda[i,8]) # firecrest

N[i,9] ~ dpois(lambda[i,9]) # goldcrest

N[i,10] ~ dpois(lambda[i,10]) # nuthatch

N[i,11] ~ dpois(lambda[i,11]) # blackcap

N[i,12] ~ dpois(lambda[i,12]) # short-toed treecreeper

N[i,13] ~ dpois(lambda[i,13]) # treecreeper

N[i,14] ~ dpois(lambda[i,14]) # stock dove

N[i,15] ~ dpois(lambda[i,15]) # black woodpecker

N[i,16] ~ dpois(lambda[i,16]) # marsh tit

\# long-tailed tit

log(lambda[i,1]) <- beta0[1] + beta[1,1] * SA[i] + beta[1,2] * MH_ric[i] + beta[1,6] * altitude[i] + beta[1,4] * formi[i] + beta[1,3] * MH_count[i] + beta[1,5] * conifer_share[i]

+ CC_AC.phi * N[i,2] + PM_AC.phi * N[i,5] + PA_AC.phi * N[i,6] + SA_AC.phi * N[i,11] + PP_AC.phi * N[i,16] + LC_AC.phi * N[i,4]

\# blue tit

log(lambda[i,2]) <- beta0[2] + beta[2,4] * formi[i] + beta[2,5] * conifer_share[i] + beta[2,6] * altitude[i] + beta[2,3] * MH_count[i] + beta[2,2] * MH_ric[i] + beta[2,1] * SA[i]

+ SA_CC.phi * N[i,11] + PM_CC.phi * N[i,5] + BW_CC.phi * N[i,15] + SE_CC.phi * N[i,10] + PA_CC.phi * N[i,6] + CO_CC.phi * N[i,14] + LC_CC.phi * N[i,4] + DM_CC.phi * N[i,3]

\# great spotted woodpecker

log(lambda[i,3]) <- beta0[3]+ beta[3,6] * altitude[i] + beta[3,1] * SA[i] + beta[3,5] * conifer_share[i] + beta[3,2] * MH_ric[i] + beta[3,3] * MH_count[i] + beta[3,4] * formi[i]

\# crested tit

log(lambda[i,4]) <- beta0[4] + beta[4,1] * SA[i] + beta[4,4] * formi[i] + beta[4,5] * conifer_share[i] + beta[4,6] * altitude[i] + beta[4,2] * MH_ric[i] + beta[4,3] * MH_count[i]

+ DM_LC.phi * N[i,3] + SE_LC.phi * N[i,10] + BW_LC.phi * N[i,15] + CO_LC.phi * N[i,14]

\# great tit

log(lambda[i,5]) <- beta0[5] + beta[5,6] * altitude[i] + beta[5,3] * MH_count[i] + beta[5,4] * formi[i] + beta[5,2] * MH_ric[i] + beta[5,5] * conifer_share[i] + beta[5,1] * SA[i]

+ SA_PM.phi * N[i,11] + DM_PM.phi * N[i,3] + LC_PM.phi * N[i,4] + PP_PM.phi * N[i,16] + CO_PM.phi * N[i,14] + SE_PM.phi * N[i,10] + BW_PM.phi * N[i,15]

\# coal tit

log(lambda[i,6]) <- beta0[6] + beta[6,1] * SA[i] + beta[6,6] * altitude[i] + beta[6,4] * formi[i] + beta[6,5] * conifer_share[i] + beta[6,3] * MH_count[i] + beta[6,2] * MH_ric[i]

+ DM_PA.phi * N[i,3] + LC_PA.phi * N[i,4] + PM_PA.phi + N[i,5] + CO_PA.phi * N[i,14] + SE_PA.phi * N[i,10] + PP_PA.phi * N[i,16] + BW_PA.phi * N[i,15]

\# chiffchaff

log(lambda[i,7]) <- beta0[7] + beta[7,1] * SA[i] + beta[7,5] * conifer_share[i] + beta[7,4] * formi[i] + beta[7,6] * altitude[i] + beta[7,2] * MH_ric[i] + beta[7,3] * MH_count[i]

+ SA_PC.phi * N[i,11] + CC_PC.phi * N[i,2] + PM_PC.phi * N[i,5] + PA_PC.phi * N[i,6] + PP_PC.phi * N[i,16] + LC_PC.phi * N[i,4]

\# firecrest

log(lambda[i,8]) <- beta0[8] + beta[8,1] * SA[i] + beta[8,2] * MH_ric[i] + beta[8,5] * conifer_share[i] + beta[8,6] * altitude[i] + beta[8,4] * formi[i] + beta[8,3] * MH_count[i]

+ LC_RI.phi * N[i,4] + PC_RI.phi * N[i,7] + RR_RI.phi * N[i,9] + PA_RI.phi * N[i,6] + SA_RI.phi * N[i,11] + PP_RI.phi * N[i,16] + PM_RI.phi * N[i,5] + CC_RI.phi * N[i,2]

\# goldcrest

log(lambda[i,9]) <- beta0[9] + beta[9,4] * formi[i] + beta[9,5] * conifer_share[i] + beta[9,1] * SA[i] + beta[9,6] * altitude[i] + beta[9,3] * MH_count[i] + beta[9,2] * MH_ric[i]

+ PC_RR.phi * N[i,7] + LC_RR.phi * N[i,4] + SA_RR.phi * N[i,11] + PP_RR.phi * N[i,16] + PA_RR.phi * N[i,6] + PM_RR.phi * N[i,5] + CC_RR.phi * N[i,2]

\# European nuthatch

log(lambda[i,10]) <- beta0[10] + beta[10,4] * formi[i] + beta[10,5] * conifer_share[i] + beta[10,6] * altitude[i] + beta[10,3] * MH_count[i] + beta[10,2] * MH_ric[i] + beta[10,1] * SA[i]

\# blackcap

log(lambda[i,11]) <- beta0[11] + beta[11,1] * SA[i] + beta[11,4] * formi[i] + beta[11,5] * conifer_share[i] + beta[11,6] * altitude[i] + beta[11,3] * MH_count[i] + beta[11,2] * MH_ric[i]

\# short-toed treecreeper

log(lambda[i,12]) <- beta0[12] + beta[12,3] * MH_count[i] + beta[12,4] * formi[i] + beta[12,6] * altitude[i] + beta[12,2] * MH_ric[i] + beta[12,1] * SA[i] + beta[12,5] * conifer_share[i]

+ PM_CB.phi * N[i,5] + PP_CB.phi * N[i,16] + SE_CB.phi * N[i,10] + PA_CB.phi * N[i,6] + DM_CB.phi * N[i,3] + BW_CB.phi * N[i,15] + CO_CB.phi * N[i,14] + CC_CB.phi * N[i,2] + LC_CB.phi * N[i,4]

\# Eurasian treecreeper

log(lambda[i,13]) <- beta0[13] + beta[13,4] * formi[i] + beta[13,3] * MH_count[i] + beta[13,6] * altitude[i] + beta[13,5] * conifer_share[i] + beta[13,1] * SA[i] + beta[13,2] * MH_ric[i]

+ CC_CF.phi * N[i,2] + LC_CF.phi * N[i,4] + BW_CF.phi * N[i,15] + PM_CF.phi * N[i,5] + SE_CF.phi * N[i,10] + DM_CF.phi * N[i,3] + PP_CF.phi * N[i,16] + PA_CF.phi * N[i,6] + CO_CF.phi * N[i,14]

\# stock dove

log(lambda[i,14]) <- beta0[14] + beta[14,4] * formi[i] + beta[14,5] * conifer_share[i] + beta[14,6] * altitude[i] + beta[14,2] * MH_ric[i] + beta[14,3] * MH_count[i] + beta[14,1] * SA[i]

\# black woodpecker

log(lambda[i,15]) <- beta0[15] + beta[15,5] * conifer_share[i] + beta[15,6] * altitude[i] + beta[15,4] * formi[i] + beta[15,2] * MH_ric[i] + beta[15,3] * MH_count[i] + beta[15,1] * SA[i]

\# marsh tit

log(lambda[i,16]) <- beta0[16] + beta[16,5] * conifer_share[i] + beta[16,2] * MH_ric[i] + beta[16,6] * altitude[i] + beta[16,1] * SA[i] + beta[16,4] * formi[i] + beta[16,3] * MH_count[i]

+ SA_PP.phi * N[i,11] + BW_PP.phi * N[i,15] + SE_PP.phi * N[i,10] + CO_PP.phi * N[i,14] + DM_PP.phi * N[i,3] + LC_PP.phi * N[i,4]

}

\# Observation model for replicated counts

for (i in 1:nsite){

for (j in 1:nrep){

Yc[i,j,1] ~ dbin(p[i,j,1], N[i,1])

Yc[i,j,2] ~ dbin(p[i,j,2], N[i,2])

Yc[i,j,3] ~ dbin(p[i,j,3], N[i,3])

Yc[i,j,4] ~ dbin(p[i,j,4], N[i,4])

Yc[i,j,5] ~ dbin(p[i,j,5], N[i,5])

Yc[i,j,6] ~ dbin(p[i,j,6], N[i,6])

Yc[i,j,7] ~ dbin(p[i,j,7], N[i,7])

Yc[i,j,8] ~ dbin(p[i,j,8], N[i,8])

Yc[i,j,9] ~ dbin(p[i,j,6], N[i,9])

Yc[i,j,10] ~ dbin(p[i,j,10], N[i,10])

Yc[i,j,11] ~ dbin(p[i,j,11], N[i,11])

Yc[i,j,12] ~ dbin(p[i,j,12], N[i,12])

Yc[i,j,13] ~ dbin(p[i,j,13], N[i,13])

Yc[i,j,14] ~ dbin(p[i,j,14], N[i,14])

Yc[i,j,15] ~ dbin(p[i,j,15], N[i,15])

Yc[i,j,16] ~ dbin(p[i,j,16], N[i,16])

logit(p[i,j,1]) <- alpha0[1] + alpha[1,1] * DAT[i,j] + alpha[1,2] * DUR[i,j]

logit(p[i,j,2]) <- alpha0[2] + alpha[2,1] * DAT[i,j] + alpha[2,2] * DUR[i,j]

logit(p[i,j,3]) <- alpha0[3] + alpha[3,1] * DAT[i,j] + alpha[3,2] * DUR[i,j]

logit(p[i,j,4]) <- alpha0[4] + alpha[4,1] * DAT[i,j] + alpha[4,2] * DUR[i,j]

logit(p[i,j,5]) <- alpha0[5] + alpha[5,1] * DAT[i,j] + alpha[5,2] * DUR[i,j]

logit(p[i,j,6]) <- alpha0[6] + alpha[6,1] * DAT[i,j] + alpha[6,2] * DUR[i,j]

logit(p[i,j,7]) <- alpha0[7] + alpha[7,1] * DAT[i,j] + alpha[7,2] * DUR[i,j]

logit(p[i,j,8]) <- alpha0[8] + alpha[8,1] * DAT[i,j] + alpha[8,2] * DUR[i,j]

logit(p[i,j,9]) <- alpha0[9] + alpha[9,1] * DAT[i,j] + alpha[9,2] * DUR[i,j]

logit(p[i,j,10]) <- alpha0[10] + alpha[10,1] * DAT[i,j] + alpha[10,2] * DUR[i,j]

logit(p[i,j,11]) <- alpha0[11] + alpha[11,1] * DAT[i,j] + alpha[11,2] * DUR[i,j]

logit(p[i,j,12]) <- alpha0[12] + alpha[12,1] * DAT[i,j] + alpha[12,2] * DUR[i,j]

logit(p[i,j,13]) <- alpha0[13] + alpha[13,1] * DAT[i,j] + alpha[13,2] * DUR[i,j]

logit(p[i,j,14]) <- alpha0[14] + alpha[14,1] * DAT[i,j] + alpha[14,2] * DUR[i,j]

logit(p[i,j,15]) <- alpha0[15] + alpha[15,1] * DAT[i,j] + alpha[15,2] * DUR[i,j]

logit(p[i,j,16]) <- alpha0[16] + alpha[16,1] * DAT[i,j] + alpha[16,2] * DUR[i,j]

}

}

\# end
